# Insights into the mutational burden of human induced pluripotent stem cells using an integrative omics approach

**DOI:** 10.1101/334870

**Authors:** Matteo D’Antonio, Paola Benaglio, David Jakubosky, William W. Greenwald, Hiroko Matsui, Margaret K. R. Donovan, He Li, Erin N. Smith, Agnieszka D’Antonio-Chronowska, Kelly A. Frazer

**Affiliations:** Institute for Genomic Medicine, University of California, San Diego, La Jolla, CA 92093, USA.; Biomedical Sciences Graduate Program, University of California, San Diego, La Jolla, CA 92093, USA.; Department of Biomedical Informatics, University of California, San Diego, La Jolla, CA 92093, USA.; Bioinformatics and Systems Biology, University of California San Diego, La Jolla, CA 92093, USA.; Department of Pediatrics and Rady Children’s Hospital, University of California San Diego, La Jolla, CA 92093, USA.

## Abstract

To understand the mutational burden of human induced pluripotent stem cells (iPSCs), we whole genome sequenced 18 fibroblast-derived iPSC lines and identified different classes of somatic mutations based on structure, origin and frequency. Copy number alterations affected 295 kb in each sample and strongly impacted gene expression. UV-damage mutations were present in ~45% of the iPSCs and accounted for most of the observed heterogeneity in mutation rates across lines. Subclonal mutations (not present in all iPSCs within a line) composed 10% of point mutations, and compared with clonal variants, showed an enrichment in active promoters and increased association with altered gene expression. Our study shows that, by combining WGS, transcriptome and epigenome data, we can understand the mutational burden of each iPSC line on an individual basis and suggests that this information could be used to prioritize iPSC lines for models of specific human diseases and/or transplantation therapy.

## Introduction

Somatic mutations in induced pluripotent stem cell (iPSCs) have been previously analyzed using a variety of approaches (Bhutani et al., 2016; Cheng et al., 2012; Gore et al., 2011; Laurent et al., 2011; Lo Sardo et al., 2017), however, a more complete understanding of mutational burden in iPSCs could increase their utility as a model system for human disease as well as for transplantation therapy. The iPSC reprogramming process involves clonal selection and somatic alterations present in the parental cell of origin or that arise during reprogramming may be under selection (Stratton et al., 2009; Torkamani et al., 2009). Previous studies that examined the genomic integrity of iPSCs have mainly used SNP arrays (International Stem Cell et al., 2011; Laurent et al., 2011; Panopoulos et al., 2017; Taapken et al., 2011), but this approach only allows the detection of relatively large copy-number alterations (CNAs, > 50 kb). A number of studies using exome or whole-genome sequencing (WGS) technologies have shown that somatic single nucleotide variants (SNVs), and small insertion and deletion (indel) mutations in iPSC lines, are predominantly derived from the parental cell rather than arising during the reprogramming process (Cheng et al., 2012; Gore et al., 2011; Kwon et al., 2017; Lo Sardo et al., 2017; Rouhani et al., 2016). Of note, the functional impact of somatic CNAs, SNVs and indels has not yet been examined in detail. Therefore, it is still unknown how to identify which somatic alterations – including those derived from the parental cell of origin as well as those that arose during reprogramming – influence molecular phenotypes in iPSCs, and hence may have an impact on the utility of iPSC-derived tissues as an experimental model of human disease and/or influence their safety for transplantation therapy.

Characterizing the somatic mutational landscape of iPSCs can be facilitated by taking advantage of the methods developed to analyze the whole-genome sequences of cancer genomes (Alexandrov et al., 2013a; International Cancer Genome et al., 2010; Nik-Zainal et al., 2016). Indeed, the analyses of cancer and iPSC genomes have several goals in common, including: 1) identification of somatic variants; 2) characterization of somatic variant function; 3) identification of the set of somatic variants under selective pressure; and 4) characterization of the subclonal nature of mutations. These cancer genomic methods compare the whole-genome sequences of tumor and a matched blood sample and are directly transferable to the analysis of somatic mutations in iPSC lines and other tissues types. A study utilizing this approach showed that in normal skin cells, which are often used to derive iPSCs, many cancer genes harbor somatic mutations that were likely caused by ultraviolet (UV) light exposure (Martincorena et al., 2015). Additionally, a study on human embryonic stem cells (Merkle et al., 2017) found that ~20% of analyzed cell lines had deleterious subclonal mutations, including some known cancer drivers in *TP53*. We hypothesize that due to their different origins, clonal and subclonal mutations have been under different selective pressures. The clonal mutations occurred in skin fibroblasts, and as mutations in somatic cells are under high selective pressure (Polak et al., 2014; Schuster-Bockler and Lehner, 2012), only neutral mutations tend to be retained. Conversely, because subclonal mutations occurred in cell culture during or after reprogramming, they have been under selective pressure for considerably less time, and thus are expected to be more likely to alter molecular phenotypes in the iPSCs and iPSC-derived cell types. Therefore, to fully understand the mutational burden of iPSCs, it is important to examine the distributions and functions of both clonal and subclonal mutations.

Many iPSC lines included in large databanks were derived from skin fibroblasts (Kilpinen et al., 2017; Panopoulos et al., 2017), and it has been estimated that up to one third of human skin cells carry UV-associated mutations in cancer genes (Martincorena et al., 2015). However, the extent to which skin fibroblast-derived iPSCs harbor UV-associated mutations has not yet been investigated. One key question is whether different iPSC lines carry similar number of mutations associated with UV damage or whether the mutational burden varies greatly across lines. Additionally, the functional impact of UV-associated mutations is important to understand, and how the utility of iPSC lines carrying large numbers of such mutations may be affected.

Here, we used deep whole-genome sequencing data (>50X average coverage) of 18 skin fibroblast-derived iPSC lines in the iPSCORE resource (Panopoulos et al., 2017) to investigate the distribution and functional impact of somatic variants, including both point mutations and larger copy-number alterations. We compared the average somatic mutational load of the 18 iPSC lines and show that it is comparable to that observed in adult stem cells, but less than what has been observed in adult tumors. We observed high variability in the number of point and indel mutations per iPSC line, which can be explained by UV exposure of the parental fibroblast cell. To assess the potential functional effects of the somatic mutations we used chromatin state information and RNA-seq data and showed that the vast majority of point mutations are in genomic regions associated with repressed chromatin and do not affect gene expression; however, compared with clonal variants, subclonal SNVs showed an enrichment in active promoters and increased association with altered gene expression. Subclonal point mutations showed a constant allelic fraction during early and late passages, as well as during differentiation into iPSC-derived cardiomyocytes, suggesting that the iPSC subclones carrying these variants do not substantially evolve in culture over time. In the 18 iPSCs we detected 255 copy number alterations which altered on average 295 kb per line and strongly impacted the expression of the genes that they overlapped. Our study demonstrates that annotating somatic mutations based on origin, structure, frequency, and the chromatin state in which they occur, enables one to predict their influence on molecular phenotypes, such as gene expression, in iPSCs and derived cell types.

## Results

### Selection and whole genome sequencing of 18 iPSC lines

We generated deep whole genome sequence data (WGS) (median read depth 50.8, range 39.8-64.6) of 18 iPSC lines previously shown to be pluripotent and to have high genomic integrity (no or low numbers of somatic CNAs) using high-throughput PluriTest-RNAseq and genotyping arrays, respectively (Panopoulos et al., 2017). The iPSC lines were derived from 18 subjects chosen to represent five different ethnicities, a range in donor ages (18 to 59 years), both sexes (12 females and 6 males) (Figure S1A; Table S1), and that are part of the 273 participants in the iPSCORE resource (many of whom have iPSC lines publicly available) (DeBoever et al., 2017; Panopoulos et al., 2017). Twelve of the 18 subjects were members of three families (one extended family and two quartets – including one with a set of identical twins, Figure S1B), and six subjects were singletons (unrelated to anyone else in this study).

### Identification of somatic mutations

To detect and characterize somatic mutations in the WGS’ of the 18 iPSC lines, we applied two established bioinformatics approaches (Mutect (Cibulskis et al., 2013) and Strelka (Saunders et al., 2012)) for analyzing cancer genome sequencing data, treating the iPSCs as “tumor” and the matched blood as “normal”. To obtain a stringent call set, we retained only somatic mutations detected using both methods and showing an allelic frequency greater than 10% (Cibulskis et al., 2013; Weinhold et al., 2014). In the genomes of the 18 iPSCs, we found 49,388 somatic mutations: 44,441 single nucleotide variants (SNVs), 2,171 dinucleotide variants (DNVs), 2,170 small deletions, and 606 small insertions (1-51 nucleotides) (Table S2). Of the 44,441 somatic SNVs, only 2,480 (5.6%) were annotated as variants in dbSNP, indicating that the somatic mutations were not misidentified inherited variants. The number of somatic mutations per iPSC line was highly variable, ranging from 958 to 7,027, corresponding to an average mutation rate of 0.88 mutations/Mb (range: 0.31 to 2.27 mutations/Mb) (Figure 1A). We found a significantly higher mutation rate (p = 8.6 × 10^−6^ Mann-Whitney U test) than previous studies examining WGS of iPSCs derived from fibroblasts (Bhutani et al., 2016) and bone marrow (Cheng et al., 2012) which respectively identified an average mutation rate of 0.25 mutations/Mb and 0.47 mutations/Mb (Figure 1B). This difference was most likely due to the fact that these previous studies compared the WGS of the iPSC lines to the parental cell population, and thus most somatic mutations present in the parental cell were excluded from the analysis; here we instead compared the WGS of the skin fibroblast-derived iPSC lines to blood DNA. We did not observe a significant association between the number of somatic mutations and the donor’s age (r = 0.107, p-value = 0.671), ethnicity (ANOVA p-value = 0.718), or gender (ANOVA p-value = 0.751). Of note, members of the same family displayed large differences in the number of somatic mutations, for instance, between the two identical twins (subjects 3_1 and 3_2 Figure 1A), there was a four-fold difference. These data show that iPSC lines have a greater number of mutations than indicated by previous studies, and suggest that the heterogeneity in mutation rates across iPSC lines is heavily influenced by a factor(s) other than donor age, ethnicity, gender or genetic background.

**Figure 1:**
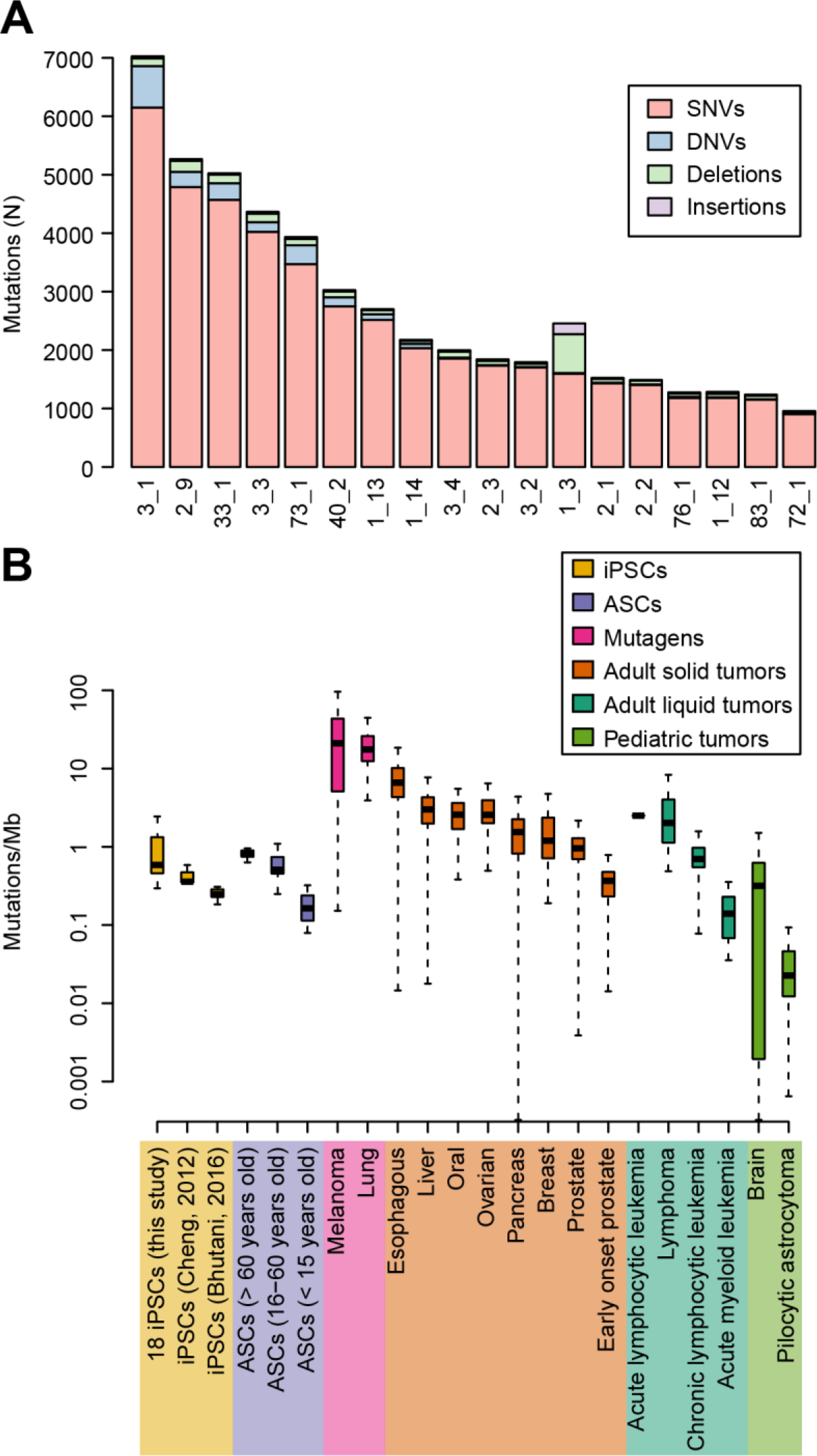
Frequency of somatic mutations in iPSCs. (**A**) Number of somatic mutations per iPSC line, divided into four types: SNVs, DNVs, small insertions and deletions. The family and subject IDs are shown (for example, 3_1 = family 3 and subject 1). (**B**) Boxplots showing the mutation rate (including SNVs, DNVs and small insertions and deletions) in the 18 iPSCORE iPSCs, in iPSCs from two previous studies (Bhutani et al., 2016; Cheng et al., 2012), adult stem cells (ASCs) (Blokzijl et al., 2016) and in 16 different cancer types (Alexandrov et al., 2013a; Berger et al., 2012). Tumors are divided into mutagens,adult solid tumors, liquid and pediatric as in Vogelstein et al. (Vogelstein et al., 2013). See also Figure S1, Figure S3, Table S1, Table S2, Table S3, Table S4.

To identify somatic copy number alterations (CNAs) with high confidence, we used Genome STRiP (Handsaker et al., 2015). CNAs were classified as somatic if: 1) they were present in the iPSCs but not in the matched blood genome; and 2) they were singleton (present only in the iPSC line and not in any of the other 255 iPSCORE WGS, including 236 blood and 19 fibroblasts). Across the 18 iPSCs, we detected 255 somatic CNAs (82 duplications and 173 deletions) which in total affected ~295 kb of sequence (0.01% of the genome) in each sample (Table S2, Table S3). The CNAs were distributed across the genome and had a median length of 1,564 bp (range 999 bp to 49 Mb). To examine the quality of our somatic CNA calls from WGS data, we compared them with somatic CNAs called from the 18 iPSCs using Illumina HumanCoreExome arrays (Panopoulos et al., 2017). From the array data, we identified 17 somatic CNAs with a median length of 259 kb (range 110 kb to 54 Mb), of which 13 (76.5%) overlapped somatic CNAs in our WGS call set (Table S4). Thus, the set of somatic CNAs identified from the WGS had high sensitivity and included hundreds of CNAs that were below the resolution afforded by SNP array analysis. Overall, we detected 255 CNAs (the majority had lengths below the resolution afforded by SNP array analysis) which altered on average 295 kb in the 18 iPSC lines.

### Comparison of iPSC mutational load to adult stem cells and tumors

To better understand the mutational load in the genomes of iPSC lines derived from skin fibroblasts, we compared the number of somatic mutations that we observed in the 18 iPSCs, to the mutation rates of adult stem cells (ASCs) and 16 different tumor types (up to 679 tumors from each type were analyzed) (Alexandrov et al., 2013a; Berger et al., 2012; International Cancer Genome et al., 2010; Nik-Zainal et al., 2016). We observed similar mutations rates between iPSCs and ASCs (Blokzijl et al., 2016) from middle age and older adult subjects (p = 0.342 and p = 0.2292, Mann-Whitney U test, respectively for subjects between 16 and 60 years old and for subjects older than 60 years), but higher mutation rates in iPSCs compared to ASCs from younger subjects (p = 2.3 × 10^−7^, Mann-Whitney U test, for subjects 15 years old or younger). We found the mutation rate in the iPSCs was substantially lower than that observed in melanoma and lung adenocarcinoma, which involve potent mutagens (ultraviolet light and cigarette smoke, respectively) in their pathogenesis (Vogelstein et al., 2013). Of note, melanoma, had significantly more somatic SNVs, DNVs, and indel mutations than the iPSCs in this study (29 times: 25.5 mutations/Mb versus 0.88, p = 6.6 × 10^−12^, Mann-Whitney U test, Figure 1B) (Berger et al., 2012). The 18 iPSCs also had a lower somatic mutation rate than most adult solid and liquid tumors (with the exception of prostate, early-onset prostate, and chronic lymphocytic leukemia), but at a higher rate than pediatric tumors (Figure 1B). These data show that the iPSCs have mutation rates comparable to stem cells but lower than most tumors obtained from individuals in the same age groups as the study subjects; however, the iPSCs have higher mutation rates than stem cells and tumors obtained from younger individuals.

### Evidence of UV damage in the skin-derived iPSC genomes

To examine the origin of the somatic mutations in the iPSCs and detect differences between lines, we compared their mutational landscapes with 30 mutational signatures derived from more than 10,000 genomes and exomes in 40 tumor types (Alexandrov et al., 2013a; Alexandrov et al., 2013b). We divided all SNV mutations into six classes of base substitutions (C>A, C>G, C>T, T>A, T>C, T>G), extracted the surrounding sequence contexts for each class and compared with the 30 mutational signatures. We found that the 18 iPSCs lines were associated with two distinct sets of mutational landscapes (Figure 2A-C, Figure S2): 1) eight samples, corresponding to the lines with the largest number of mutations (Table S2), are strongly correlated with Signature 2 (prevalence of C>T transitions, likely due to the AID/APOBEC family of cytidine deaminases), Signature 7 (caused by UV exposure and predominantly found in skin cancers), Signature 11 (associated with melanoma, likely due to the presence of alkylating agents) and Signature 30 (prevalence of C>T transitions, of unknown origin); and 2) ten samples are strongly correlated with mutational signatures associated with a high prevalence of C>A transversions (Signatures 4, 8, 10, 16, 18 and 29) and with Signature 5, which is found ubiquitously in all cancer types but is not associated with any known process (Helleday et al., 2014). These results suggest that the parental cells used to reprogram the eight iPSC lines with highest amounts of mutations had been subjected to UV damage.

In the 18 iPSC lines we investigated the prevalence of CNAs, C>T SNVs and CC>TT DNVs, of which the latter two mutation classes are typical of melanoma and are known to be caused by UV damage (Brash, 2015; Brash et al., 1991; Gartner et al., 2013; Greenman et al., 2007). The eight most mutated iPSCs had significantly more CNAs (22.3 CNAs/samples on average) than the other ten samples (7.5 CNAs/samples on average, p = 0.032, Mann-Whitney U test), but the total amount of the genome involved in CNAs was not higher (295 kb and 292 kb, respectively, p = 0.75, Mann-Whitney U test, Table S2) (Figure 2C). We found that the eight most mutated iPSCs had a relatively high incidence of C>T SNVs (Figure 2D) and CC>TT DNVs (Figure 2E, Table S5), and that the number of these mutations in a given line were highly correlated with the total number of mutations (C>T: r = 0.98, CC>TT: r = 0.91, Figure 2C). The other DNV classes together correspond to 0.7% of all somatic mutations, while CC>TT accounted for 4.8% in the eight most mutated iPSC lines (range 3.0 to 9.2%). Overall, these results show that 45% of the iPSC lines were derived from parental fibroblast cells harboring UV damage, that these iPSCs have significantly more CNAs, C>T SNVs and CC>TT DNVs, and mutational signatures similarly to those found in melanoma.

**Figure 2:**
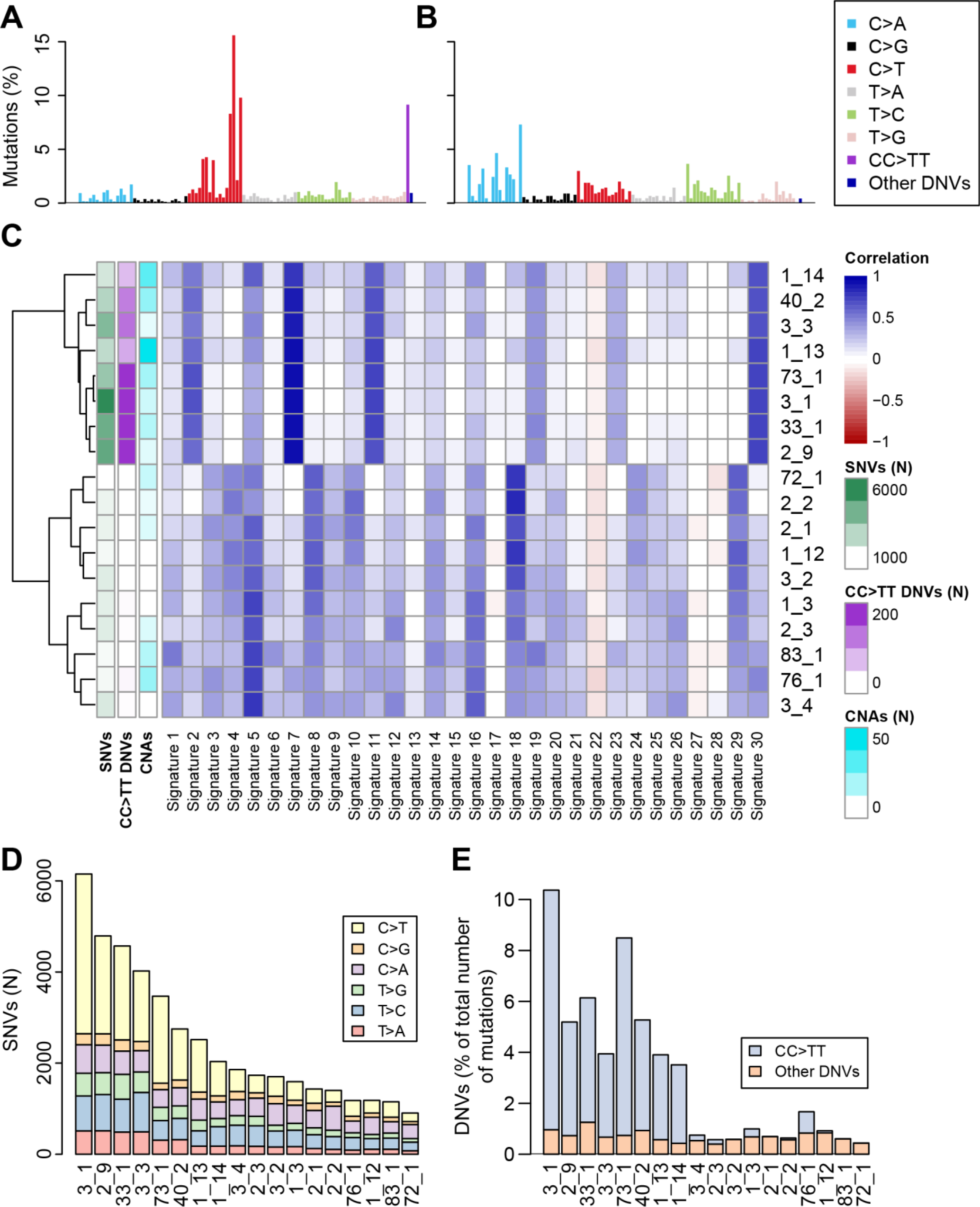
Evidence of UV damage in iPSCs. (**A**, **B**) Plots showing the percentage of mutations in each of the 96 substitution classes defined by the substitution type and one base sequence context immediately 3’ and 5’ to the mutated base. (**A**) Example iPSC line with UV damage (sample 3_1); (**B**) Example of iPSC line without UV damage (72_1). (**C**) Heatmap showing the correlation between the mutational profiles of the 18 iPSC lines and 30 mutational signatures derived from 40 tumor types (Alexandrov et al., 2013b). The number of SNVs, CC>TT DNVs and CNAs are shown. (**D**) Distribution of SNV types in each of the 18 iPSC lines. (**E**) The fraction of all mutations in each of the 18 iPSC lines that are DNVs. Eight iPSC lines have a high incidence of CC>TT DNVs (4.8% of all point and indel mutations in the eight most mutated iPSC lines), while the number of the other DNVs is low and constant in all 18 iPSCs (0.4-1.2% of all somatic mutations). See also Figure S2, Table S5.

### Functional impact of clonal and subclonal mutations on gene structure

While clonal mutations present in all cells of an iPSC line are likely to be derived from the parental cell, those that are subclonal and only present in a fraction of the cells, must have arisen during the reprogramming process or subsequent culturing. Based on the frequency of the mutated allele in the iPSC line, we divided the somatic mutations into three classes: 1) clonal (frequency of mutated allele between 30% and 80%); 2) subclonal (frequency of mutated allele between 10% and 30%); and 3) hemizygous (frequency of mutated allele > 80%) (Figure 3). We found the majority of somatic SNVs (39,100; 88.0%) and DNVs (1,602; 88.1% of CC>TT, and 306; 86.7% of the other DNVs) were clonal (Table S2). There were 4,738 subclonal SNVs (10.7% of all SNVs, range=4.0-26.8% per iPSC line, Table S2) and 263 subclonal DNVs (12.1 % of all DNVs). Additionally, we identified 603 hemizygous SNVs and DNVs (1.29% of the total number of somatic mutations), located on sex chromosomes in male samples or in regions where their sister chromosome contained a large deletion. These results indicate that ~11% of all somatic mutations are subclonal and based on their frequency (10% to 30%) likely arose within the first few cellular divisions after reprogramming of the parental cell.

**Figure 3:**
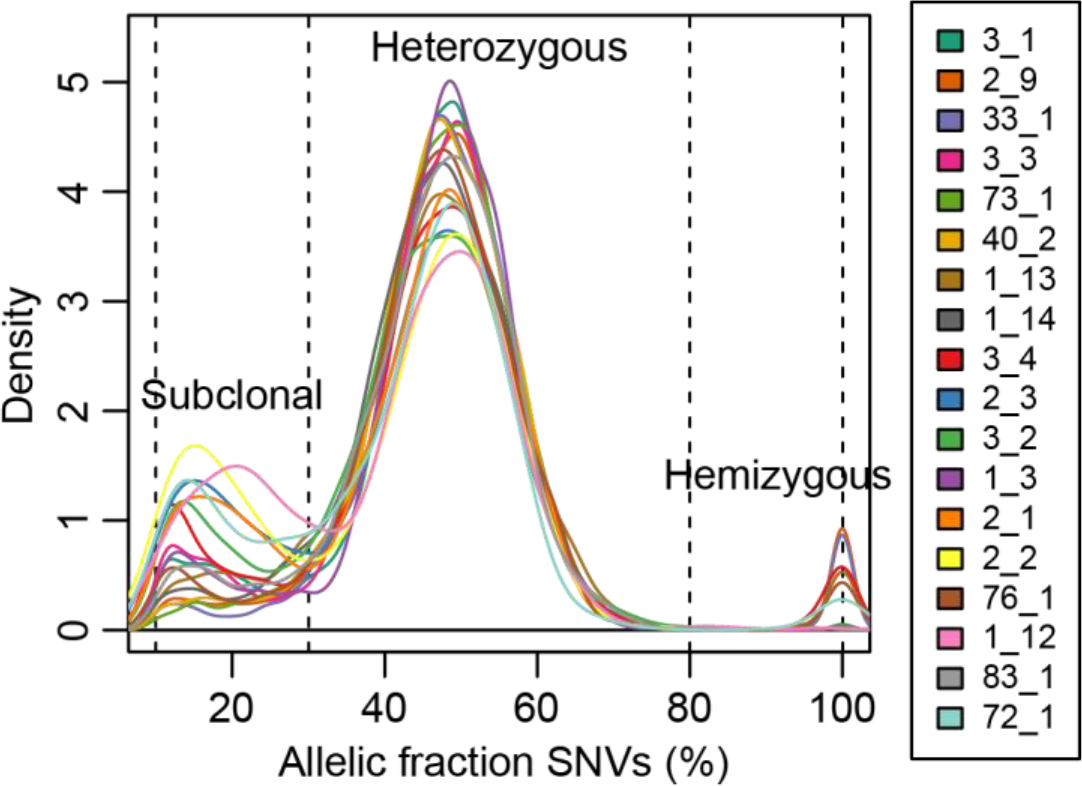
Clonal and subclonal SNVs. Density plot of the allelic fraction distribution of SNVs for each iPSC line, showing that the vast majority of mutations are heterozygous. The peak at 20% allelic fraction corresponds to subclonal variants that occurred during early stages of reprogramming. The peak at 100% corresponds to hemizygous mutations. See also Figure S1.

We examined if the different origins of clonal and subclonal mutations resulted in their having different functional impacts on gene structure. We grouped somatic mutations into four classes: 1) clonal C>T SNVs; 2) other clonal SNVs; 3) clonal CC>TT DNVs; and 4) subclonal SNVs. We considered clonal C>T SNVs separately from the other clonal SNVs and only the CC>TT DNVs, because these two classes are likely caused by UV exposure in the parental cell of origin (Brash, 2015; Brash et al., 1991; Gartner et al., 2013; Greenman et al., 2007). We annotated the variants using SnpEff (Cingolani et al., 2012), which divides the mutations into four groups based on their predicted functional impact on gene structure (Table S6): 1) no impact (mutations occurring in intergenic regions or intronic regions that do not affect splice sites); 2) low impact (mutations affecting UTRs, transcription factor binding sites, non-coding exons and synonymous mutations); 3) moderate impact (missense, splice-site and in-frame indels); and 4) high impact (nonsense and frameshift). In total, we found 504 genes affected by low, moderate, or high impact mutations (186, 287 and 31 genes, respectively, Table S7), and an average of 21 moderate or high impact mutations per iPSC line – substantially more than previous studies, which identified 2-12 non-synonymous mutations per iPSC line (Bhutani et al., 2016; Cheng et al., 2012; Gore et al., 2011) (Table S6). These findings are likely because we identified iPSC somatic mutations by comparison with blood DNA rather than with DNA from the parental tissue of origin, as was done in the prior studies (Bhutani et al., 2016; Cheng et al., 2012; Gore et al., 2011). While all four classes of somatic mutations had similar fractions of no impact (~97%), moderate impact (~0.8%) and high impact (~0.04%) variants, subclonal SNVs had a significantly greater fraction of low impact variants compared with clonal SNVs (2.5% versus 1.5%, p = 1.5 × 10^−7^, Fisher’s exact test, Figure 4A). Overall these analyses suggest that iPSC lines carry ~2 times more detrimental coding mutations than estimated by previous studies, and show that while the majority of SNVs and indels have no predicted functional impact on gene structure, there was an enrichment of low impact mutations in subclonal SNVs compared with clonal SNVs.

**Figure 4:**
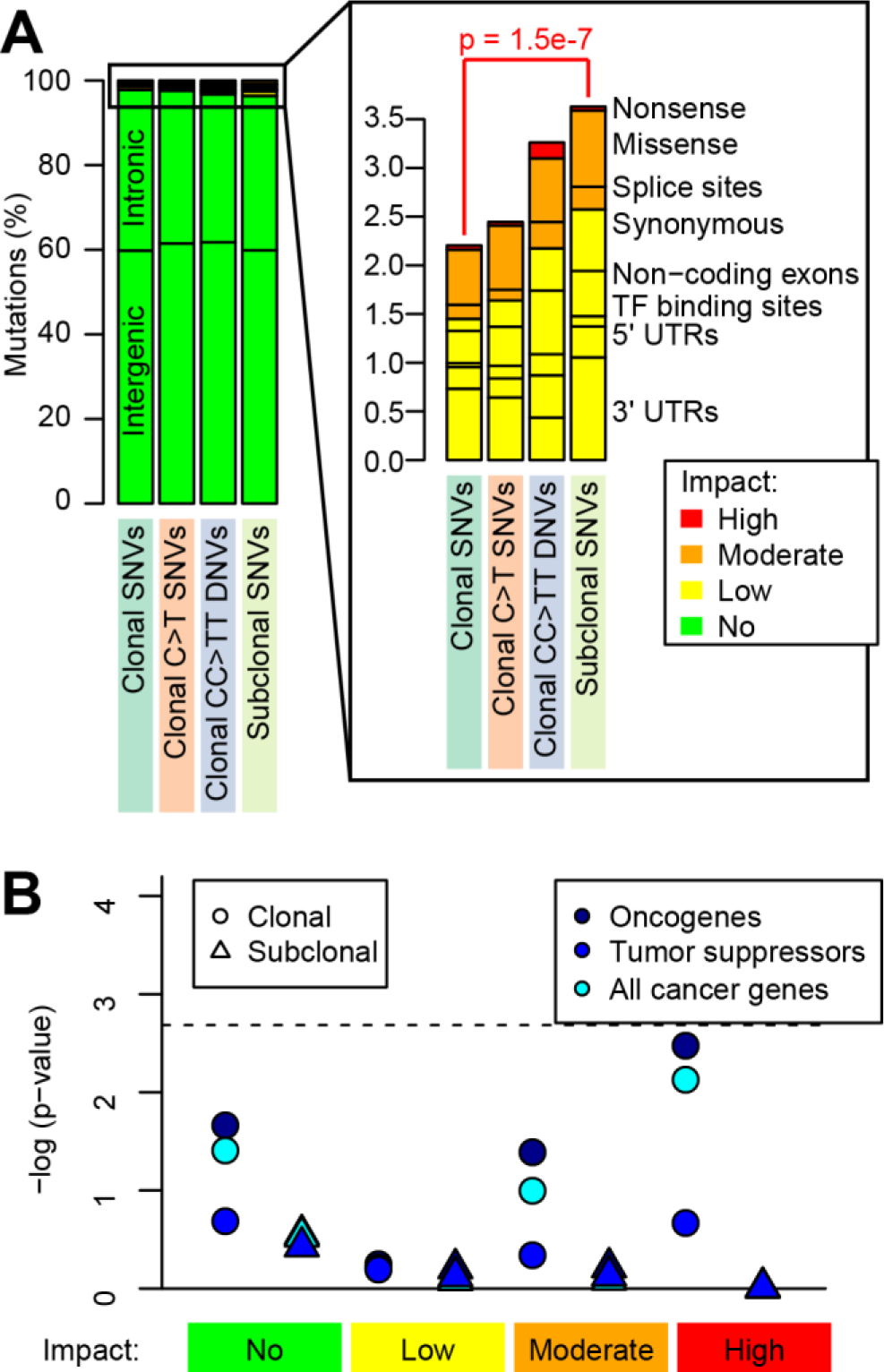
Functional characterization of somatic SNVs and DNVs. (**A**) For the four somatic mutation classes, the variants are grouped based on impact determined via SnpEff (Cingolani et al., 2012). P-value shows the enrichment of low impact mutations in subclonal SNVs compared with clonal SNVs (Fisher’s exact test). (**B**) Enrichment analysis for clonal (the union of the three classes of clonal variants) and subclonal variants in cancer genes. The horizontal dashed line shows the p-value threshold of significance for Bonferroni correction (FDR < 0.2). See also Table S6, Table S7.

As it has been reported that embryonic stem cells may harbor subclonal mutations in cancer genes (Merkle et al., 2017), we investigated whether the iPSCs showed enrichment for clonal (combined the three classes: clonal SNVs, clonal C>T SNVs and clonal CC>TT DNVs) or subclonal SNVs and indels in cancer genes. We intersected cancer-associated genes using the Cancer Gene Census (Forbes et al., 2015) (110 oncogenes and 141 tumor suppressors) with 504 genes that we identified as carrying either high, moderate, low, or no impact mutations in one or more of the 18 iPSC lines (Table S7). Of the 504 genes carrying mutations, only 13 were cancer genes (eleven carried clonal and two carried subclonal mutations), which was not significantly different than expected by chance (Figure 4B). Of note, eight iPSC lines carried clonal non-synonymous mutations in one or more known oncogenes (*COL1A1, IL6ST, JUN*, and *NUP214)*, tumor suppressors (*BLM, CYLD, FANCA, POLE* and *STAG2*), and *IKZF1*, which behaves as either an oncogene or a tumor suppressor in different tumor types, and two lines carried subclonal non-synonymous mutations in two oncogenes (*DDX6* and *PDE4DIP*) (Table S7). These results indicate that neither clonal nor subclonal somatic mutations are enriched in cancer genes; but by chance ~50% of the iPSC lines carry moderate or high impact mutations in known cancer genes.

### Clonal and subclonal mutations are associated with different chromatin states

We investigated the distributions of clonal and subclonal somatic mutations with respect to active and repressed chromatin states (Ernst and Kellis, 2012) in 22 stem cell lines from the Roadmap and ENCODE consortia (Neph et al., 2012; Roadmap Epigenomics et al., 2015). As expected (Yoshihara et al., 2017), we observed that overall mutations were more likely than expected to occur in repressed chromatin regions (defined as heterochromatin, polycomb repressed and quiescent chromatin) and less likely to occur in active chromatin (transcribed regions and active promoters, Figure 5A). A direct comparison of the distribution between the clonal SNVs and C>T SNVs across the 15 chromatin states showed that they were highly correlated (r = 0.961) (Figure 5A); both mutation classes were strongly enriched in repressive chromatin states (heterochromatin, weak repressed polycomb and quiescent chromatin regions) and strongly depleted in active chromatin states [transcribed genes, enhancers, transcriptional start sites (TSS)]. However, clonal C>T SNVs were significantly more likely to occur in heterochromatin than other clonal SNVs (p = 6.41 × 10^−5^, t-test, Bonferroni correction for FDR; Figure 5B, Table S8). The distributions of clonal SNVs and CC>TT DNVs were slightly more weakly correlated (r = 0.850, Figure 5A) and CC>TT DNVs were less likely to occur in repressive chromatin states, including heterochromatin and quiescent chromatin regions (p = 1.77 × 10^−10^ and p = 1.48 × 10^−6^, respectively) and more likely to be present in transcribed genes (p = 3.27 × 10^−6^), active TSS (p = 1.00 × 10^−12^) or in enhancers (p = 3.69 × 10^−8^). Compared with clonal SNVs, subclonal SNVs demonstrated significantly different associations with chromatin marks (r = 0.596): they were not enriched in quiescent regions (p = 1.43 × 10^−12^) and not strongly depleted in active TSS (p = 9.36 × 10^−15^), weakly transcribed genes (p = 1.87 × 10^−14^), strongly transcribed genes (p = 3.88 × 10^−6^), 5’ and 3’ transcribed regions (p = 6.01 × 10^−5^). These data demonstrate that the four mutational classes are associated with different chromatin states, and that subclonal SNVs are the most likely to be located in functional genomic regions.

**Figure 5:**
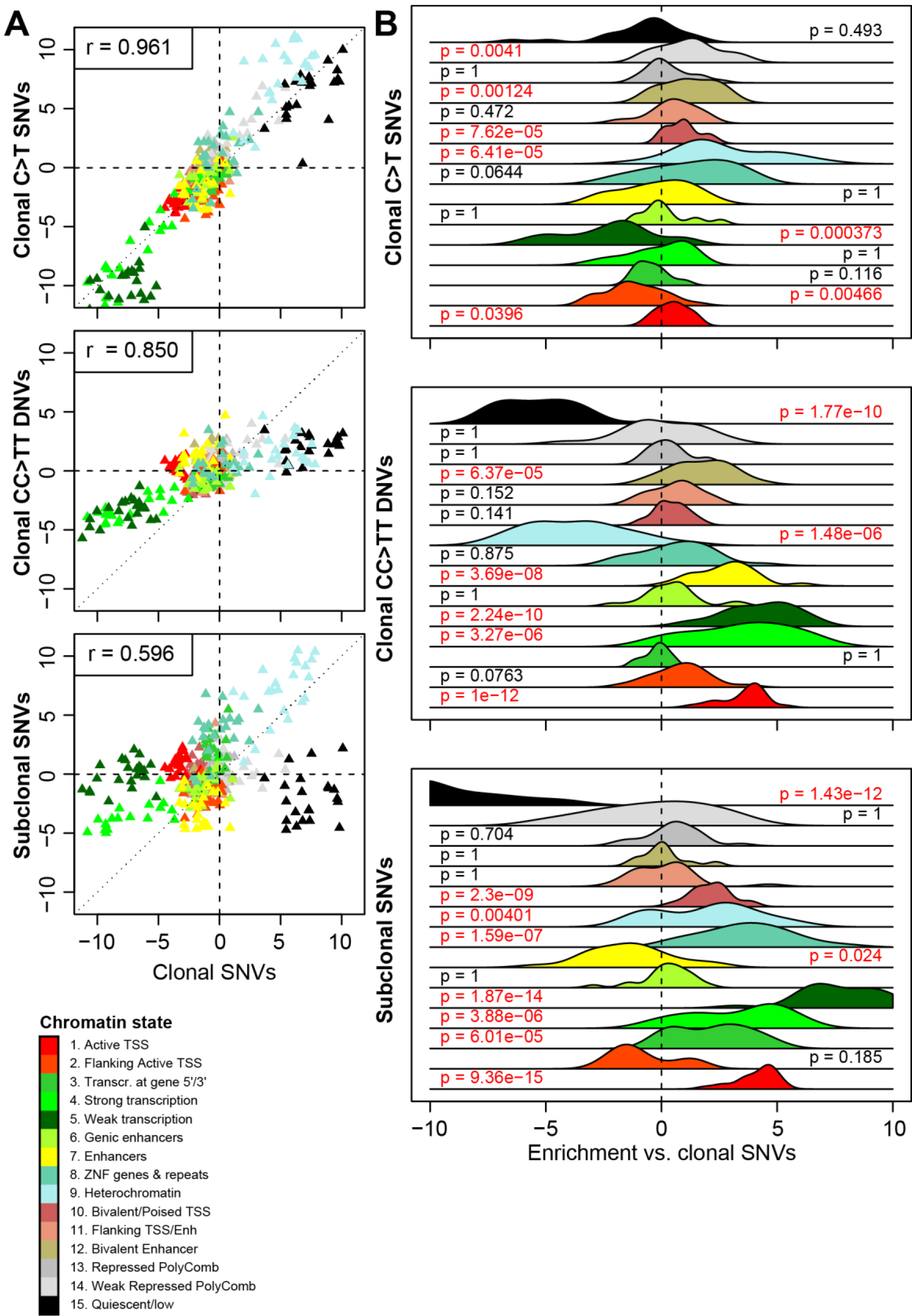
Associations between somatic mutations and chromatin states. (**A**) Scatterplots showing the enrichment Z-score of each chromatin mark in the 22 stem cell lines from Roadmap and ENCODE for clonal SNVs (X axis in all three plots) compared with clonal C>T SNVs (Y axis), clonal CC>TT DNVs (Y axis) and subclonal SNVs (Y axis). (**B**) The enrichment of clonal C>T SNVs, CC>TT DNVs and 22subclonal SNVs mutations compared with clonal SNVs in each of 15 ChromHMM chromatin states for 22 stem cell lines are shown as a plot of the distribution of Z-scores differences (each class of mutations – clonal SNVs). P-values were calculated for each chromatin mark by comparing the Z-score distributions between clonal SNVs and the other classes of mutations across the 22 stem cell lines using paired t-test and FDR-adjusted using Bonferroni’s method. See also Table S8.

### Subclonal mutations and CNAs result in altered gene expression

To examine if the different associations between the four mutation classes and chromatin states could result in different effects on cellular molecular phenotypes, we investigated their associations with aberrant gene expression using an approach developed by the GTEx Consortium to determine the effects of rare genetic variation on gene expression (Li et al., 2017). For each gene, we quantile-normalized its expression levels across 222 iPSC lines included in the iPSCORE cohort (DeBoever et al., 2017), we found its closest mutation (<500 kb from its TSS) and determined its normalized expression level in the mutated sample (Table S9). We next compared the distribution of normalized expression levels between genes with clonal SNVs and genes with either subclonal SNVs, clonal CC>TT DNVs or clonal C>T SNVs (Figure 6A). We observed that subclonal SNVs were more likely to be associated with aberrantly expressed genes than clonal SNVs (p = 0.019, Fisher’s exact test), whereas CC>TT DNVs (p = 0.036) and C>T SNVs (p = 0.0029) were less likely. An example of a functional subclonal SNV includes a mutation present at 29% frequency located 14 kb upstream of *MYCL* associated with a large decrease in expression level (TPM is reduced from 20.8 to 8.6, Figure 6B). *MYCL is* a member of the myelocytomatosis oncogene (MYC) family involved in cell proliferation and death, is required for iPSC reprogramming and for dendritic cell differentiation (Hatton et al., 1996; Kc et al., 2014; Nakagawa et al., 2010). Of note, the closest mutation to a gene was a clonal SNV which occurred in the promoter of *IRF2* (131 bp upstream) and resulted in a 75% increase (Z-score = 2.15) of its expression level (TPM = 12.3; mean TPM across all iPSC lines = 7.0, Figure 6C). This gene is a member of the interferon regulatory transcription factor (IRF) family, and its overexpression has been shown to have pro-oncogenic activity (Cui et al., 2012; Harada et al., 1993). These results show that while subclonal mutations were more likely to be associated with alterations of gene expression levels than clonal mutations, regulatory mutations included in all mutational classes may alter expression levels of genes that are involved in differentiation and/or cancer.

**Figure 6:**
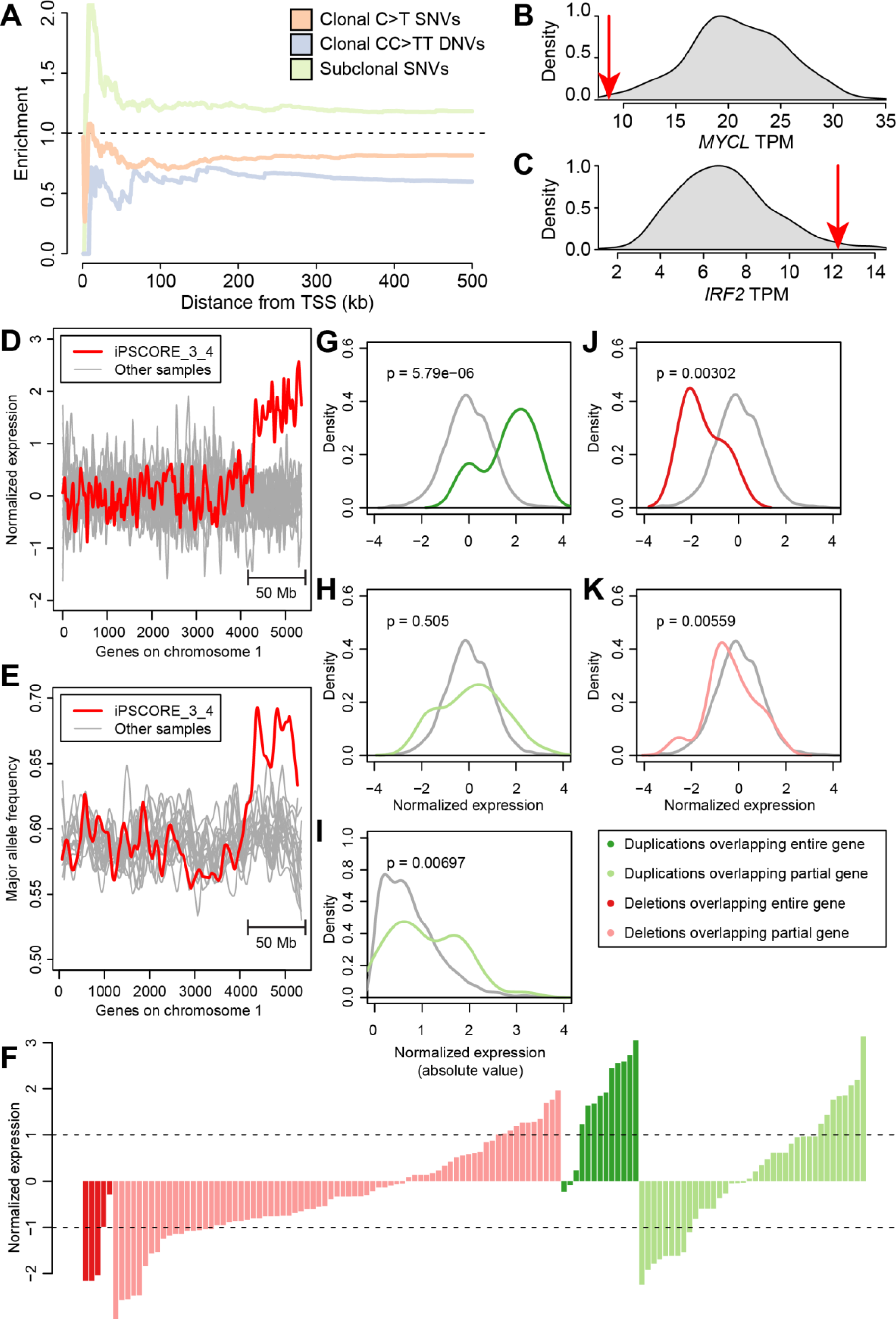
Differential effects of somatic mutations on gene expression. (**A**) Relative enrichment for clonal C>T SNVs, clonal CC>TT DNVs and subclonal SNVs compared with clonal SNVs. Enrichment was calculated as ratio between the fraction of differentially expressed (Z-score >2 or < −2) mutated genes in each mutational class and clonal SNVs in 1-kb bins. (**B, C**) Density plots showing the TPM distribution across 222 iPSC lines in the iPSCORE collection for (**B**) *MYCL* and (**C**) *IRF2*. Red arrows represent the expression levels in the samples harboring the mutation. (**D**,**E**) Effects of a 50 Mb duplication of chromosome 1 on gene expression. The expression levels of genes in iPSCORE_3_4 are shown in red, while the expression of these genes in the other iPSCs are shown as gray lines. The X axis is ordered according to the location of the genes on chromosome 1 based on genomic coordinate. (**D**) Normalized expression levels, calculated as Z-scores of all genes on chromosome 1. (**E**) Major allele expression frequency for all genes on chromosome 1, as calculated by MBASED(Mayba et al., 2014), the large CNA results in a major allele expression frequency of ~0.67, consistent with the presence of three copies of the q arm. (**F**) Normalized expression levels of genes overlapping the other 254 CNAs divided into four categories, based on whether they are a duplication or deletion and their overlap with genes: 1) deletions that overlap an entire gene (dark red); 2) deletions that partially overlap a gene (light red); 3) duplications that overlap an entire gene (dark green); and 4) duplications that partially overlap a gene (light green). (**G-K**) Comparison between the mean expression levels of genes overlapping CNAs (colored as in panel **F**) and the mean expression levels of the same genes in iPSC lines that do not carry the CNA (gray); (**G**) genes completely included within a large duplication; (**H**) genes partially overlapped by a large duplication; (**I**) absolute value of the normalized expression levels of genes partially overlapped by a duplication; (**J**) genes completely included within a large deletion; (**K**) genes partially overlapped by a large deletion. Gene expression was normalized to have mean = 0 and standard deviation = 1 for each gene. See also Table S9, Table S10.

We next investigated the functional impact of the 255 CNAs in the 18 iPSC lines on gene expression (Table S10). We initially examined the effect of the largest CNA identified (a duplication of the distal 50 Mb of chromosome 1q in iPSCORE_3_4). As expected based on gene dosage, we determined that the expression of genes in iPSCORE_3_4 overlapping this CNA were on average 1.6 standard deviations higher than the mean average of the same genes in the other iPSC lines (p = 2.1 × 10^−91^, t test) (Figure 6D). Additionally, they were more likely to show allele specific expression (ASE) compared to the other iPSCs (p = 3.2 × 10^−32^, t test) and to the other chromosome 1 genes in iPSCORE_3_4 outside of the duplication (2.3 × 10^−3^, t test) (Figure 6E). Congruent with these findings, 10 of the 13 genes whose sequence was fully included in CNA duplications were overexpressed (p = 5.8 × 10^−6^, t test, Figure 6F,G). While the mean expression levels of 38 genes that only partially overlapped an CNA duplication did not display significant overexpression or downregulation (p = 0.51, Figure 6H), we observed that nine of these genes had a normalized expression level below −1 SD and eight above 1 SD (Figure 6F), suggesting that several genes in this group may indeed be aberrantly expressed. Therefore, we investigated the absolute gene expression levels of these 38 genes that partially overlapped a CNA duplication, and found that their absolute expression levels were significantly higher than expected, confirming that genes that partially overlapped a CNA duplication could have altered expression (p = 0.00697, t test, Figure 6I). Genes affected by somatic CNA deletions were downregulated, irrespective of whether the deletion overlapped the entire gene (n = 5; p = 3.0 × 10^−3^, t test) or only part of the gene (n = 78; p = 5.6 × 10^−3^, t test, Figure 6J,K). These results show that of the 134 genes that are expressed in iPSCs and overlap somatic CNAs, 48 (35.8%) had significantly altered expression in iPSC lines.

### iPSCs carrying subclonal SNVs do not evolve

To understand the extent to which subclonal mutations in iPSCs evolve, we examined changes in their allele frequencies during culturing and differentiation into cardiomyocytes by analyzing RNA-seq data as recently described by Merkle et al. (Merkle et al., 2017). For each of the four iPSC lines of family 2 (iPSCORE_2_1, iPSCORE_2_2, iPSCORE_2_3 and iPSCORE_2_9; Figure S1, Table S11) between six and twenty-one independent RNA sequencing datasets were generated at varying passages (P12 to P25) and time points (day 2, 5, 9 and 15) during cardiomyocyte differentiation (Panopoulos et al., 2017). We were able to analyze the allelic frequency of 146 coding mutations, of which 30 were subclonal, of which only four (2.4%) (one subclonal, three clonal with allelic frequency < 40%) significantly changed allelic fraction (Bonferroni-adjusted p-value < 0.1, Cochran-Armitage test for trend). Thus, the vast majority of subclonal SNVs tended to maintain a constant allelic fraction at all time points (Figure 7), suggesting that different subclones were in equilibrium and did not substantially evolve between passage 12 and later passages, or throughout differentiation. This consistency in allele frequencies across up to seven distinct time points and more than 50 cell divisions suggests that most iPSC subclonal mutations are stable in culture (i.e. not under strong selective pressure), and thus not evolving over time.

**Figure 7:**
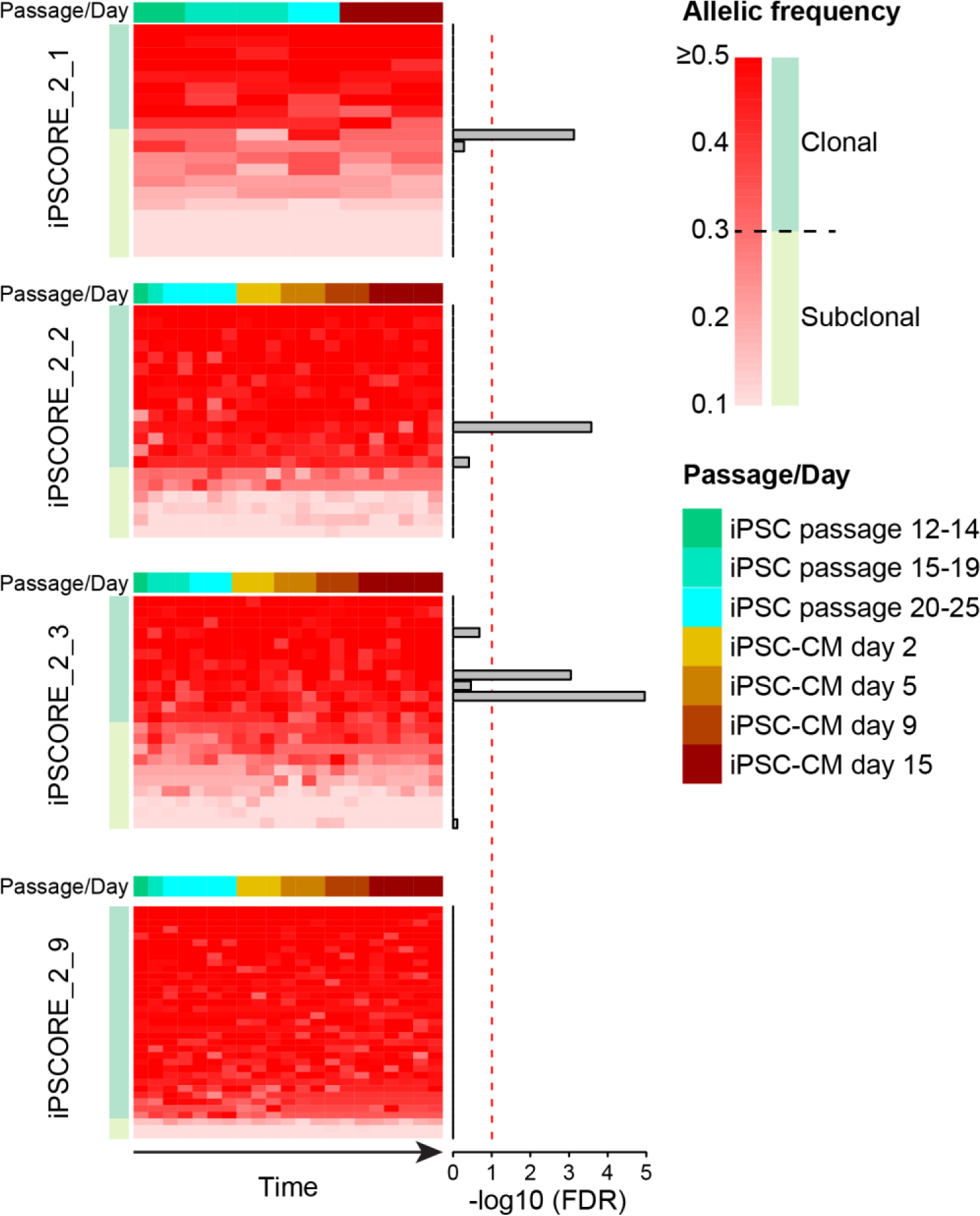
Subclonal evolution of somatic mutations. Heatmaps showing the allelic frequency for all point and indel mutations in four subjects with RNA-seq at multiple iPSC passages and during iPSC-CM differentiation. Each row in the heatmaps represents a somatic mutation in a transcribed region (146 mutations in total); each column represents a different time point. For each individual, mutations were sorted from the highest allelic fraction to the lowest. Barplots next to each mutation represent Cochran-Armitage p-values adjusted for FDR (Bonferroni’s method), showing that allelic frequency changes only for four mutations across different passages and differentiation. See also Table S11.

## Discussion

In this study, we used deep whole-genome sequencing data (>50X average coverage) of 18 iPSC lines to investigate the distribution and functional impact of somatic variants, including both point mutations and larger copy number alterations. Because we identified iPSC somatic mutations by comparison with blood DNA, rather than with DNA from the parental tissue of origin, we found that the mutational burden in iPSCs (including both point mutations and structural variants) is greater than it was previously reported (Bhutani et al., 2016; Cheng et al., 2012; Rouhani et al., 2016). The use of deep WGS data also enabled us to also detect 20 times more CNAs at higher resolution (1 kb) than previous studies have reported (International Stem Cell et al., 2011; Laurent et al., 2011; Taapken et al., 2011). We did not observe significant associations between the number of mutations and donor age, ethnicity, gender or genetic background; however, we discovered that UV-associated mutations are a major contributor to the heterogeneity in mutation rates across iPSC lines. Of note, we show that the mutational load in iPSCs is comparable to what is observed in adult stem cells, and hence somatic mutations may have similar effects on molecular phenotypes in both cell types. Our findings were consistent with previous studies showing that mutations present in iPSCs are underrepresented in gene bodies and in genomic regions associated with open chromatin (Yoshihara et al., 2017), and that the origin of somatic mutations influences their functional features (Rouhani et al., 2016). Overall, our analyses not only provided insights about the functional impact of two previously investigated classes of somatic mutations (clonal SNVs and CNAs) but also resulted in the discovery and functional characterization of two previously undescribed classes of somatic mutations (UV damage and subclonal SNVs) in iPSCs.

Mutational signatures associated with UV damage (high prevalence of clonal C>T SNVs and clonal CC>TT DNVs) were found in ~45% of the iPSC lines and were likely derived from the parental skin-fibroblast cell. The iPSC lines displaying UV damage mutational signatures had significantly more CNAs, but the total amount of the genome involved in CNAs was not higher, suggesting that UV damage typically does not result in large chromosomal alterations. We observed that clonal CC>TT DNVs were more likely to occur in open chromatin than other clonal mutations (clonal SNVs); however, we also observed that clonal C>T and clonal CC>TT DNVs tend to impact gene expression less than other clonal mutations. These two observations suggest that UV-associated mutations (even those in functional chromatin states) are less likely to affect molecular phenotypes than other clonal mutations. Although mutations caused by UV damage appear to be largely neutral, for any given iPSC line harboring a UV damage mutational signature, due to the sheer number of mutations some are likely to strongly impact a molecular phenotype.

Approximately 10% of all somatic mutations in the iPSC lines were subclonal, which suggests that a single line may contain multiple subclones with highly heterogeneous genetic backgrounds. The allelic fraction of subclonal mutations remained constant between early and later iPSC passages and during differentiation into cardiomyocytes, suggesting that they were in an equilibrium state under culturing conditions. Additionally, subclonal mutations were more likely to occur within active chromatin regions (in particular promoters and transcribed regions) than clonal mutations. Furthermore, subclonal mutations had significantly stronger positive effects on gene expression than clonal SNV mutations. The functional characteristics of subclonal mutations suggest that they have been under less negative selection than clonal mutations (likely due to their origin during or shortly after reprogramming). Of note, since subclonal mutations are present only in a fraction of the iPS cells in a line (20% to 60% of cells), their effects on gene expression within a specific cell were likely stronger than what we observed.

Until recently, CNAs have been difficult to call from WGS data, and therefore most studies investigating these alterations in iPSC lines have analyzed CNAs called from array data. Having high-coverage WGS data (>50X), we were able to detect 255 CNAs in the 18 iPSC lines, the majority of which had lengths below the resolution afforded by SNP array analysis, that altered on average 295 kb in each line. We investigated the effects of somatic CNAs on gene expression and found that more than one third of the genes overlapped by CNAs had altered expression, in a fashion similar to rare inherited CNAs (Chiang et al., 2016; DeBoever et al., 2017). These findings show that one can predict the effect of a CNA on the expression of overlapping genes in iPSC lines based on whether the alteration is a duplication (upregulate) or deletion (downregulate).

In summary, we extensively characterized somatic mutations in iPSCs based on their structures and origins and found that different mutational classes display different properties. Although some mutational classes were more likely to be functional than others, we detected hundreds to thousands of mutations per iPSC line, and showed that mutations in all classes may be associated with altered molecular phenotypes. While the effects of some mutations could be predicted based on their structure and genomic location, most had to be paired with an RNA-seq sample from the same iPSC line to fully characterize their effect on gene expression. It is likely that some of the somatic mutations that did not affect gene expression in the iPSCs may have affects in specific differentiated tissues. For instance, mutations in a cardiac specific transcription factor, such as *NKX2-5*, would potentially have effects on phenotypes in iPSC-derived cardiomyocytes but not in iPSC-derived neurons or in the iPSCs themselves. In conclusion, our study shows that, by combining WGS, transcriptome and epigenome data, we can understand the mutational burden of each iPSC line on an individual basis and can use this information to prioritize the use of specific iPSC lines for modeling human diseases and/or transplantation therapy.

## Experimental Procedures

### Samples and iPSC reprogramming

The 18 iPSC lines analyzed in this study are part of the iPSCORE resource (Panopoulos et al., 2017). 273 individuals were recruited into the iPSCORE study and whole genome sequencing of their blood or skin fibroblast DNA (254 DNA samples isolated from blood and 19 DNA samples isolated from skin fibroblasts) was conducted as previously described (DeBoever et al., 2017; Jakubosky et al., In preparation). The iPSCORE iPSC lines were systematically derived as described in Panopoulos et al. (Panopoulos et al., 2017).

### Whole genome sequencing

WGS was performed as previously described (DeBoever et al., 2017). The reads were aligned to human genome hg19 with decoy sequences (Genomes Project et al., 2015) using BWA-MEM with default parameters (Li and Durbin, 2009). Duplicate reads were marked using Biobambam2 (Tischler and Leonard, 2014), and reads were sorted by genomic coordinate using Sambamba (Tarasov et al., 2015) in BAM format. The 18 iPSC WGS data (and the previously generated matched blood WGS) were high quality, having 4-20% duplicates and a minimum of 700M reads after duplicate removal (Figure S3).

### Somatic SNVs and indel calling

We used Mutect (Cibulskis et al., 2013) to detect somatic SNVs and Strelka (Saunders et al., 2012) to detect SNVs and small indels that were present in DNA isolated from the 18 iPSC lines but not in the DNA isolated from matched blood. Results from the two variant callers were intersected and only SNVs called with both methods were considered as valid somatic mutations. DNVs were identified by merging two SNVs with distance = 1 bp between each other.

### Somatic CNA detection

We used the population level read-depth and split-read caller Genome STRiP (svtoolkit 2.00.1611) to discover and genotype CNAs (duplications, deletions and multi-allelic CNAs) in the 18 iPSCs and their matched blood genomes (Handsaker et al., 2015). We ran Genome STRiP using the suggested settings for high coverage genomes. We considered the CNAs as somatic mutations if they: 1) were present in the iPSC line but not the matched blood genome and; 2) they were singleton, i.e. present only in one iPSC genome and not present in any of the additional 256 genomes without matched iPSCs (Jakubosky et al., In preparation).

### Somatic mutations in adult stem cells and cancer

Somatic mutations in adult stem cells (ASCs) from three different tissue types (liver, small intestine and colon) derived from 45 subjects were obtained from Blokzijl et al. (Blokzijl et al., 2016). The number of somatic mutations in each subject for each tumor were obtained from three collections: 1) 507 tumors (four tumor types) from Alexandrov et al. (Alexandrov et al., 2013a); 2) 25 melanomas from Berger et al. (Berger et al., 2012); and 3) 3,011 tumors (11 tumor types) from the International Cancer Genomics Consortium (ICGC) (Alexandrov et al., 2013a; International Cancer Genome et al., 2010; Nik-Zainal et al., 2016).

### Detection of associations between mutations and chromatin states

The whole genome was divided into 200-bp bins and the bins with similar sequence characteristics (and thus similar mutation rates) were clustered together using four covariates (D’Antonio et al., 2017; Lawrence et al., 2013): 1) DNA replication timing; 2) open vs. closed chromatin status; 3) GC content; and 4) gene density in the 500 kb surrounding each bin. The values of all covariates were normalized to have mean = 0 and standard deviation = 1. Normalized covariate values were used to cluster all 200-bp bins in each chromosome using k-means clustering, where k was selected to have on average 200 bins in each cluster. A BED file was created for each cluster. Each somatic variant (including SNVs and small indels) was assigned to its 200 bp bin and its position was permuted 100 times within all the sequences in its cluster. The number of variants associated with each chromatin mark was determined in all permutations, then mean and standard deviation were calculated. Enrichment for each mutation class in each chromatin state was determined on a per tissue basis as Z-scores against the 100 permutations by subtracting from the observed value its corresponding mean across the 100 permutations and dividing by the standard deviation.

### RNA-seq data processing and gene expression analysis

We used previously published RNA-seq data for iPSC lines at passage 12 (DeBoever et al., 2017). For each gene, TPM values were normalized using the calcNormFactors function in the preprocessCore package in R. For 18,284 genes with mean TPM >0, we calculated the distance between their TSS and their closest upstream mutation. For each gene, we determine the normalized expression value in the mutated sample, expressed in Z-scores (defined as the number of standard deviations from the mean). To determine enrichment between different mutational classes, we calculated the fraction of genes with Z-score >2 or < −2 and compared it with clonal SNVs for all mutations within 500 kb from the TSS.

### Somatic CNA impact on gene expression

To assess overlap between CNAs and genes, first, we intersected the coordinates of each CNA with the coordinates of GENCODE genes using Bedtools intersect and found 1,325 genes that overlapped the 255 CNAs, of which 1,049 overlapped the 50 Mb chromosome 1q duplication in iPSCORE_3_4 (Table S10). For each gene that overlapped a CNA, we compared its normalized expression level in the iPSC line that harbored the CNA with respect to all other lines. To analyze the large duplication on chromosome 1, allelic-bias was determined using WASP (van de Geijn et al., 2015) and allele specific expression was calculated using MBASED (Mayba et al., 2014).

### Assessing clonal evolution of iPSCs using RNA-seq

To assess evolution of somatic mutations in iPSCs, we used 71 independent RNA-seq data sets from three other iPSCORE studies (Benaglio et al., 2018; DeBoever et al., 2017; Panopoulos et al., 2017). The RNA-seq data were generated from iPSCs at varying passages ~12-25, and iPSC-derived cardiomyocytes (iPSC-CMs) at differentiation day 15 for four subjects and a time course analysis of iPSC-CM differentiation (days 2, 5 and 9) in three subjects (Table S11).

Detailed methods are provided in the Supplemental Experimental Procedures.

## Accession numbers

Phenotype, array genotypes, RNA-seq data, and whole genome sequence genotypes are available through dbGaP (dbGaP: phs000924 and phs001325). The 222 iPSC lines are available through WiCell Research Institute (www.wicell.org;NHLBINextGenCollection).

## Author contributions

KAF and MD conceived the study. MD performed data processing and computational analyses. DAJ performed CNA detection from WGS. HM performed whole genome sequencing data processing. MD, WWG and MKRD performed gene expression analysis. PB, WWG and HL performed Hi-C sample preparation and analysis. AA isolated DNA and RNA from the iPSCs. ADC oversaw the experimental procedures. ENS oversaw WGS and RNA-seq analyses. MD and KAF wrote the paper.

## Acknowledgements

This work was supported in part by a California Institute for Regenerative Medicine (CIRM) grant GC1R-06673 and NIH grants HG008118-01, HL107442-05, DK105541-03 and DK112155-01. RNA-seq were performed at the UCSD IGM Genomics Center with support from NIH grant P30CA023100. DJ and MKRD were supported by the National Library of Medicine Training Grants T15LM011271. We thank Bing Ren, Siddarth Selvaraj and Anthony Schmitt for generating the Hi-C data. Whole genome sequencing was performed at Human Longevity, Inc.

## Declaration of interests

The authors declare no competing interests.

